# Virulence in a *Pseudomonas syringae* Strain with a Small Repertoire of Predicted Effectors

**DOI:** 10.1101/000869

**Authors:** Kevin L. Hockett, Marc T. Nishimura, Erick Karlsrud, Kevin Dougherty, David A. Baltrus

## Abstract

Both type III effector proteins and non-ribosomal peptide toxins play important roles for *Pseudomonas syringae* pathogenicity in host plants, but whether and how these virulence pathways interact to promote infection remains unclear. Genomic evidence from one clade of *P. syringae* suggests a tradeoff between the total number of type III effector proteins and presence of syringomycin, syringopeptin, and syringolin A toxins. Here we report the complete genome sequence from *P. syringae* CC1557, which contains the lowest number of known type III effectors to date and has also acquired genes similar to sequences encoding syringomycin pathways from other strains. We demonstrate that this strain is pathogenic on *Nicotiana benthamiana* and that both the type III secretion system and a new type III effector family, *hopBJ1*, contribute to virulence. We further demonstrate that virulence activity of HopBJ1 is dependent on similar catalytic sites as the *E. coli* CNF1 toxin. Taken together, our results provide additional support for a negative correlation between type III effector repertoires and the potential to produce syringomycin-like toxins while also highlighting how genomic synteny and bioinformatics can be used to identify and characterize novel virulence proteins.

## Introduction

*Pseudomonas syringae*, a bacterial phytopathogen of many crop species, utilizes a diverse arsenal of virulence factors to infect host plants (Hirano and Upper, 2000; O’Brien et al., 2011). Much of the previous research on pathogenicity in *P. syringae* has focused on identifying and characterizing virulence factors within strains and on categorizing their presence across the species (Baltrus et al., 2011; Hwang et al., 2005; Lindeberg et al., 2012; O’Brien et al., 2011). Although this accumulation of knowledge has enabled tests of protein function during infection, many questions remain as to how virulence pathways interact and evolve at a systems level (Baltrus et al., 2011; Cunnac et al., 2011; Hwang et al., 2005). In the absence of direct tests, identification of similar genomic trends across independent lineages provides strong support for the presence of such interactions over evolutionary time scales.

One of the main contributors to virulence within *P. syringae* is a type III secretion system (T3SS) (O’Brien et al., 2011). The T3SS encodes a structure that translocates bacterial effector proteins (T3E) into host cells to disrupt host physiological pathways and enable successful infection (Lindeberg et al., 2012). *P. syringae* genomes typically contain about 10-40 total T3Es, the exact actions of which depend on the totality of effector repertoires as well as host genotype (Baltrus et al., 2011; Lindeberg et al., 2012; O’Brien et al., 2011). Another main contributor to pathogenesis within many *P. syringae* strains is non-ribosomal peptide (NRP) toxin pathways. NRP pathways encode proteins that act as an assembly line, elongating and decorating peptide chains involved in pathogenicity (Bender et al., 1999). NRP pathways are often not conserved between closely related strains, but even if present, can differ in their outputs based on nucleotide polymorphisms (Baltrus et al., 2011; Bender et al., 1999; Hwang et al., 2005).

Multiple reports have suggested functional overlap or phenotypic interactions between T3E and NRP pathways. For instance, phenotypic virulence functions of the toxin coronatine can be partially restored by the toxin syringolin A as well as T3Es HopZ1 and AvrB1 (Cui et al., 2010; Jiang et al., 2013; Melotto et al., 2006; Schellenberg et al., 2010). Furthermore, within MLSA group II *P. syringae* strains as defined by Sarkar and Guttman (2004), there exists a negative correlation between presence of conserved toxin pathways (syringolin A, syringopeptin, and syringomycin) and size of the T3E repertoire compared to other phylogenetic clades (Baltrus et al., 2011; Figure S1). In fact, MLSA group II strains possess the lowest number of T3E among analyzed genomes across the species. This correlation is strengthened by the observation that strains from pathovar *pisi* possess some of the largest T3E repertoires within MLSA group II and have lost all three toxin pathways (Baltrus et al., 2013). Whether this negative correlation between NRPs and T3Es reflects an unrecognized difference in disease ecology is yet to be determined, although strains from MLSA group II are thought to survive as epiphytes much better than other clades within *P. syringae* (Feil et al., 2005).

The hypothesis of a phenotypic tradeoff between T3E and NRP pathways would be bolstered by discovery of an independent clade of *P. syringae* that has gained similar toxins to the MLSA group II strains but which also possesses a reduced effector repertoire. We have analyzed the genomes of a small clade of *P. syringae* isolated as from environmental sources (Morris et al., 2008; Baltrus et al. 2014), and have found independent evidence supporting this genomic trend (Figure S1). Specifically, compared to a closely related outgroup, genomes for two strains appear to have lost T3Es while gaining a pathway which potentially encodes proteins for the production of a syringomycin-like toxin. We demonstrate that one of these strains, CC1557, can infect *Nicotiana benthamiana* and cause disease in a T3SS-dependent manner. We further show that virulence of this strain is significantly increased by the presence of one new T3E effector family, HopBJ1, which shares similar structure and catalytic residues with the CNF1 from *Escherichia coli*. Therefore, independent evidence suggests that acquisition NRP pathways correlates strongly with the loss of T3E families and further strengthens the idea of an ecological or evolutionary tradeoff between these virulence factors.

## Results

### A complete genome sequence for *P. syringae* CC1557

Using a combination of 100bp Illumina paired end reads and longer PacBio reads, we assembled a complete genome sequence for strain CC1557 (Morris et al. 2008). This genome consists of a 5,728,024 bp chromosome and a 53,629 bp plasmid (Genbank accession number AVEH02000000). According to PGAAP annotation (Angiuoli et al. 2008), this chromosome contains 5001 ORFs while the single plasmid contains an additional 67 ORFs.

### *P. syringae* CC1557 can infect and cause disease in *N. benthamiana*

*P. syringae* CC1557 was originally isolated from snow, while the closely related strain CC1466 was originally isolated from asymptomatic *Dodecantheon pulchellum*, a perennial herb (Morris et al., 2008). Using syringe inoculation under standard conditions, we have demonstrated that *P. syringae* CC1557 can grow to high population sizes in the apoplast of *N. benthamiana* after 3 days of infection (Figure 1). This rate of growth is similar to the well-established pathogenic strain for *N. benthamiana, P. syringae* pv. *syringae* B728a (Figure S2). Furthermore, these large bacterial population sizes cause disease symptoms as evidenced by the visibility of tissue collapse (Figure 3). High levels of growth and tissue collapse are both eliminated by deleting the *hrcC* gene (Figures 1 and 3), which encodes a main structural protein for the type III secretion pilus. Therefore, CC1557 virulence in *N. benthamiana* under laboratory conditions is T3SS dependent.

**Figure 1.**
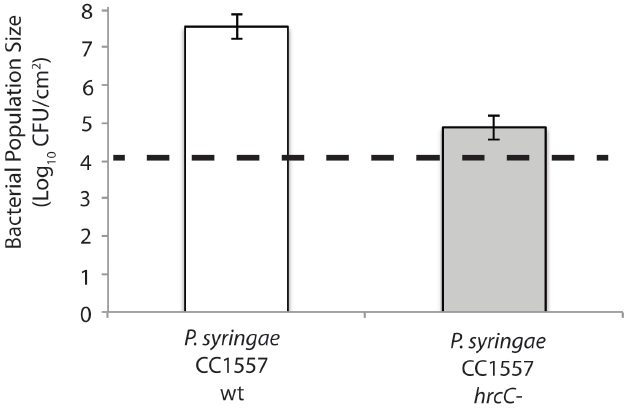
*P. syringae* CC1557 Virulence in *N. benthamiana* is Type III Secretion Dependent. **A)** Growth of wild type *P. syringae* CC1557 as well as a *hrcC* mutant was measured *in planta* three days post syringe inoculation. Measurements are based on three independent experiments with at least 4 replicates in each. Bacterial population sizes are significantly different (Tukey’s HSD, p < 0.05) between strains. Error bars represent 2 standard errors. Dashed lines indicate approximate population sizes at day 0, which were not significantly different.

**Figure 2.**
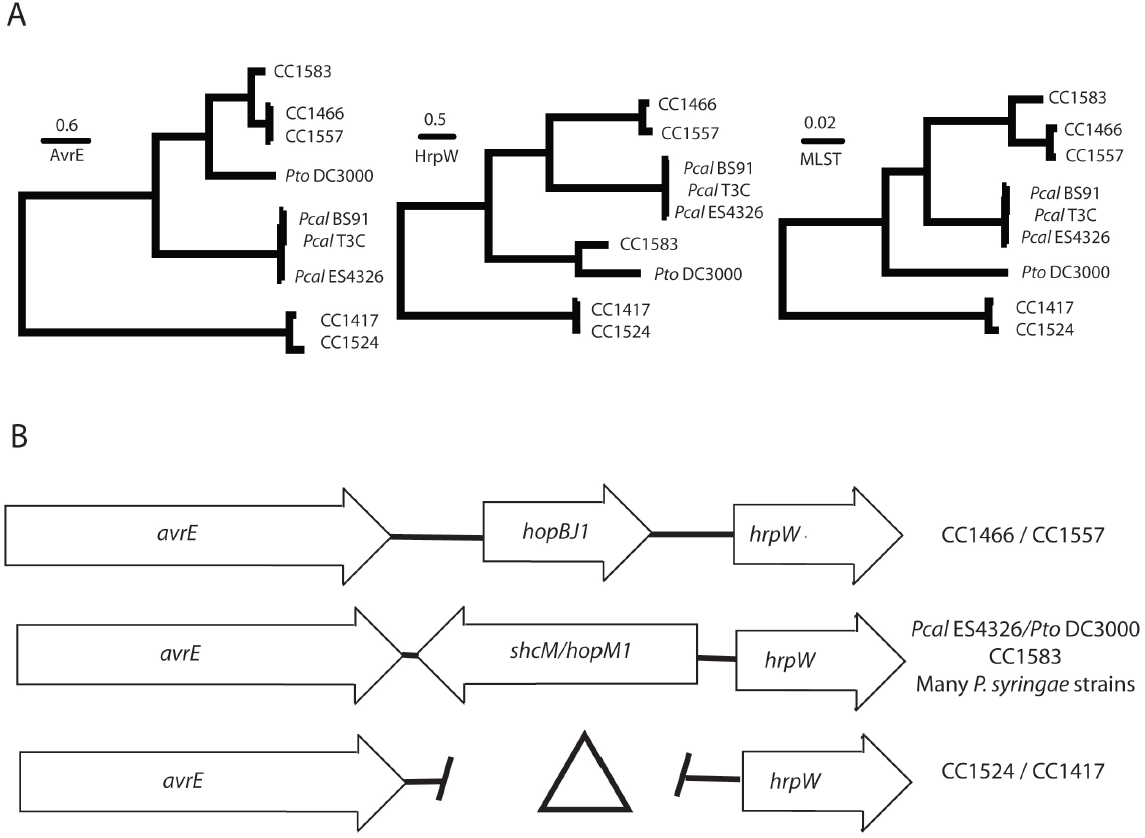
Diversity of the CEL locus across *P. syringae sensu latu*. **A)** Bayesian phylogenies for AvrE (left), HrpW (middle), and concatenated MLSA loci (right), for a subset of *P. syringae* strains. Posterior probabilities for all nodes are >0.95. Scale bars indicate number of amino acid changes. Phylogenetic patterns for both of these loci approximate relationships based on core genomes and MLSA loci, except that HrpW from *P. syringae* CC1583 clusters more closely with *P. syringae* pv. *tomato* DC3000. This suggests a recombination event at this locus within *P. syringae* CC1583. **B)** Genomic context of the CEL across *P. syringae sensu latu.* In most strains, *hopM1* is bordered by *avrE* and *hrpW.* Some *P. viridiflava* strains lack *hopM1* (Bartoli et al., 2014), while CC1557 and CC1466 have replaced this region with *hopBJ1*. Arrows in represent ORFs while the gap and triangle represent a deletion of this region within strains CC1524 and CC1417.

**Figure 3.**
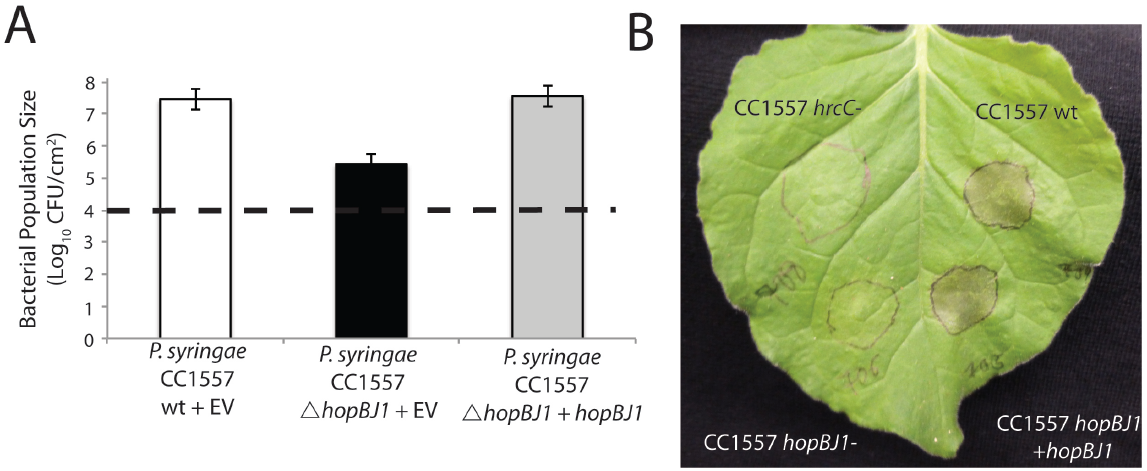
HopBJ1 is a Virulence Factor for *P. syringae* CC1557 in *N. benthamiana*. We created a deletion mutant of *hopBJ1* in *P. syringae* CC1557, and complemented this strain by expressing *hopBJ1* from a plasmid using the native promoter. Deletion of *hopBJ1* lowers growth of *P. syringae* CC1557 three days post inoculation. Both of these phenotypes are complemented by expression of wild type *hopBJ1 in trans*. Three independent experiments were performed with at least 4 replicates in each treatment. Letters represent statistical differences within an ANOVA (Tukey’s HSD, p < 0.05). Error bars represent 2 standard errors. Dashed lines indicate approximate population sizes at day 0, which were not significantly different. **B)** Leaves from two-week old *N. Benthamiana* plants were syringe inoculated with various strains derived from *P. syringae* CC1557. Three days after inoculation, plants showed evidence of tissue collapse dependent on a functioning T3SS (compare wild type vs. *hrcC* mutant) as well as HopBJ1 (compare *hopBJ1* deletion strain vs. *hopBJ1* complement). Picture is representative of multiple trials.

#### The genomes of *P. syringae* CC1557 and CC1466 encode a reduced type III effector repertoire

We used bioinformatic methods as per Baltrus et al. 2011, to search the complete and draft genomes of *P. syringae* strains CC1557, CC1466, and the closely related strain CC1583, for known type III effector families (Baltrus et al. 2014). We found a total of four T3E shared across these three genomes, with eight T3E found only within CC1583 (Table 2). Therefore, together with HopBJ1 (see below), the genomes of CC1466 and CC1557 appear to encode a total of 5 potential T3E from known protein families. This stands in contrast to the immediate outgroup strain CC1583, which encodes a total of 12 potential T3E from known protein families.

**Table 1.**
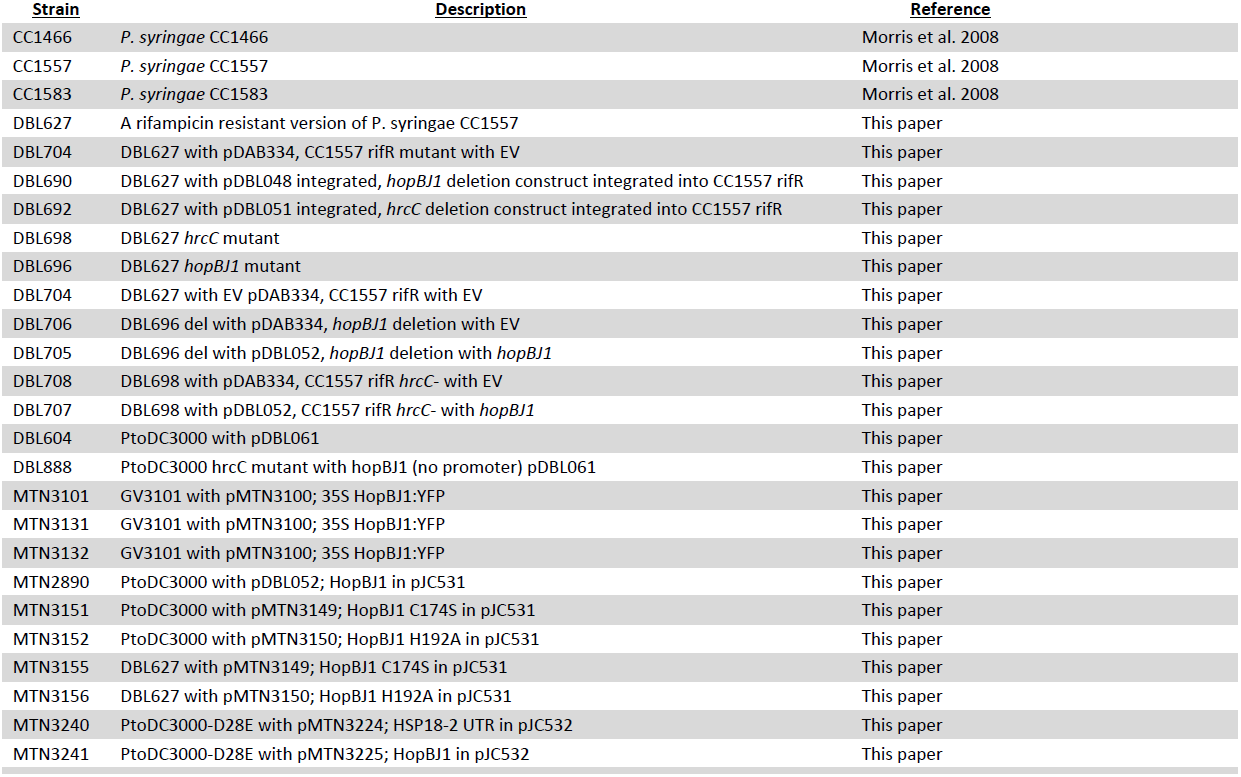
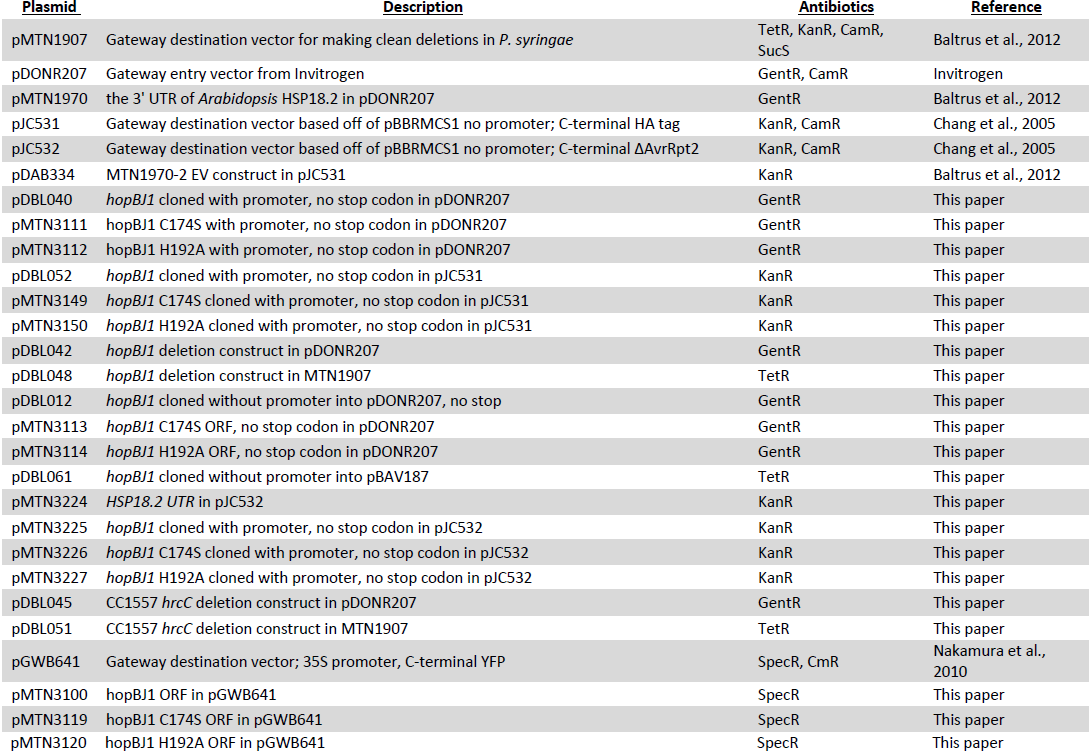
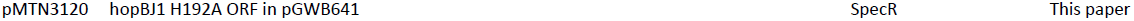
Strains and Plasmids.

| **Strain** | **Description** | **Reference** |
| --- | --- | --- |
| CC1466 | *P. syringae* CC1466 | Morris et al. 2008 |
| CC1557 | *P. syringae* CC1557 | Morris et al. 2008 |
| CC1583 | *P. syringae* CC1583 | Morris et al. 2008 |
| DBL627 | A rifampicin resistant version of *P. syringae* CC1557 | This paper |
| DBL704 | DBL627 with pDAB334, CC1557 rifR mutant with EV | This paper |
| DBL690 | DBL627 with pDBL048 integrated, *hopBJ1* deletion construct integrated into CC1557 rifR | This paper |
| DBL692 | DBL627 with pDBL051 integrated, *hrcC* deletion construct integrated into CC1557 rifR | This paper |
| DBL698 | DBL627 *hrcC* mutant | This paper |
| DBL696 | DBL627 *hopBJ1* mutant | This paper |
| DBL704 | DBL627 with EV pDAB334, CC1557 rifR with EV | This paper |
| DBL706 | DBL696 del with pDAB334, *hopBJ1* deletion with EV | This paper |
| DBL705 | DBL696 del with pDBL052, *hopBJ1* deletion with *hopBJ1* | This paper |
| DBL708 | DBL698 with pDAB334, CC1557 rifR *hrcC*- with EV | This paper |
| DBL707 | DBL698 with pDBL052, CC1557 rifR *hrcC*- with *hopBJ1* | This paper |
| DBL604 | PtoDC3000 with pDBL061 | This paper |
| DBL888 | PtoDC3000 *hrcC* mutant with *hopBJ1* (no promoter) pDBL061 | This paper |
| MTN3101 | GV3101 with pMTN3100; 35S *HopBJ1*:YFP | This paper |
| MTN3131 | GV3101 with pMTN3100; 35S *HopBJ1*:YFP | This paper |
| MTN3132 | GV3101 with pMTN3100; 35S *HopBJ1*:YFP | This paper |
| MTN2890 | PtoDC3000 with pDBL052; *HopBJ1* in pJC531 | This paper |
| MTN3151 | PtoDC3000 with pMTN3149; *HopBJ1* C174S in pJC531 | This paper |
| MTN3152 | PtoDC3000 with pMTN3150; *HopBJ1* H192A in pJC531 | This paper |
| MTN3155 | DBL627 with pMTN3149; *HopBJ1* C174S in pJC531 | This paper |
| MTN3156 | DBL627 with pMTN3150; *HopBJ1* H192A in pJC531 | This paper |
| MTN3240 | PtoDC3000-D28E with pMTN3224; HSP18-2 UTR in pJC532 | This paper |
| MTN3241 | PtoDC3000-D28E with pMTN3225; *HopBJ1* in pJC532 | This paper |

| <u>Plasmid</u> | <u>Description</u> | <u>Antibiotics</u> | <u>Reference</u> |
| --- | --- | --- | --- |
| pMTN1907 | Gateway destination vector for making clean deletions in *P. syringae* | TetR, KanR, CamR, SucS | Baltrus et al., 2012 |
| pDONR207 | Gateway entry vector from Invitrogen | GentR, CamR | Invitrogen |
| pMTN1970 | the 3' UTR of *Arabidopsis* HSP18.2 in pDONR207 | GentR | Baltrus et al., 2012 |
| pJC531 | Gateway destination vector based off of pBBRMCS1 no promoter; C-terminal HA tag | KanR, CamR | Chang et al., 2005 |
| pJC532 | Gateway destination vector based off of pBBRMCS1 no promoter; C-terminal ΔAvrRpt2 | KanR, CamR | Chang et al., 2005 |
| pDAB334 | MTN1970-2 EV construct in pJC531 | KanR | Baltrus et al., 2012 |
| pDBL040 | *hopBJ1* cloned with promoter, no stop codon in pDONR207 | GentR | This paper |
| pMTN3111 | *hopBJ1* C174S with promoter, no stop codon in pDONR207 | GentR | This paper |
| pMTN3112 | *hopBJ1* H192A with promoter, no stop codon in pDONR207 | GentR | This paper |
| pDBL052 | *hopBJ1* cloned with promoter, no stop codon in pJC531 | KanR | This paper |
| pMTN3149 | *hopBJ1* C174S cloned with promoter, no stop codon in pJC531 | KanR | This paper |
| pMTN3150 | *hopBJ1* H192A cloned with promoter, no stop codon in pJC531 | KanR | This paper |
| pDBL042 | *hopBJ1* deletion construct in pDONR207 | GentR | This paper |
| pDBL048 | *hopBJ1* deletion construct in MTN1907 | TetR | This paper |
| pDBL012 | *hopBJ1* cloned without promoter into pDONR207, no stop | GentR | This paper |
| pMTN3113 | *hopBJ1* C174S ORF, no stop codon in pDONR207 | GentR | This paper |
| pMTN3114 | *hopBJ1* H192A ORF, no stop codon in pDONR207 | GentR | This paper |
| pDBL061 | *hopBJ1* cloned without promoter into pBAV187 | TetR | This paper |
| pMTN3224 | *HSP18.2* UTR in pJC532 | KanR | This paper |
| pMTN3225 | *hopBJ1* cloned with promoter, no stop codon in pJC532 | KanR | This paper |
| pMTN3226 | *hopBJ1* C174S cloned with promoter, no stop codon in pJC532 | KanR | This paper |
| pMTN3227 | *hopBJ1* H192A cloned with promoter, no stop codon in pJC532 | KanR | This paper |
| pDBL045 | CC1557 *hrcC* deletion construct in pDONR207 | GentR | This paper |
| pDBL051 | CC1557 *hrcC* deletion construct in MTN1907 | TetR | This paper |
| pGWB641 | Gateway destination vector; 35S promoter, C-terminal YFP | SpecR, CmR | Nakamura et al., 2010 |
| pMTN3100 | *hopBJ1* ORF in pGWB641 | SpecR | This paper |
| pMTN3119 | *hopBJ1* C174S ORF in pGWB641 | SpecR | This paper |

**Table 2.** Type III Effector Family Distribution.

|  | CC1557 | CC1466 | CC1583 |
| --- | --- | --- | --- |
| *avrE* | P | P | P |
| *hopF2-1* |  |  | P |
| *hopF2-2* |  |  | P |
| *hopM1* |  |  | P |
| *hopY1* | P | P | P |
| *hopAA1* |  |  | P |
| *hopAG1* |  |  | P |
| *hopAH1* |  |  | P |
| *hopAH2* | P | P | P |
| *hopAI1* |  |  | P |
| *hopAS1* |  |  | P |
| *hopBF1* | P | P | P |
| *hopBJ1* | P | P |  |

The letter P denotes presence of this particular effector family, whereas the lack of a letter denotes absence.

#### *P. syringae* CC1466 and CC1557 have horizontally acquired pathways for production of non-ribosomal peptides that resemble syringopeptin and syringomycin

We used tBLASTn searches of the complete genome sequence of CC1557 in order to identify potential NRP toxin pathways as per Baltrus et al. 2011. This genome contains highly similar full-length tBLASTn matches to a variety of proteins involved in syringomycin synthesis: SyrB (93% Protein Sequence Identity), SyrC (99%), and SyrP (93%), data not shown. Further investigation of all complete genomes for *P. syringae* demonstrates that the region containing putative NRP-related loci in CC1557 is found in approximately the same genomic context on the chromosome as the syringomycin-syringopeptin pathways in *P. syringae* pv. *syringae* B728a (Figure S3). Moreover, the genomic context surrounding these toxin pathways is conserved throughout *P. syringae* (Figure S3). Although regions of high similarity to the syringopeptin pathway do not appear to be present within CC1557, analysis of gene annotations suggests that a second NRP pathway occurs immediately downstream of the syringomycin-like region on the chromosome of CC1557 (data not shown). While it is difficult to identify the NRP product of these CC1557 pathways by sequence alone, draft genomes sequences suggest that this region is present in CC1466 but not CC1583. The most parsimonious explanation for this phylogenetic signal is that an immediate ancestor of CC1466 and CC1557 acquired this NRP region by horizontal transfer after divergence from the CC1583 lineage.

#### A new type III effector protein family is present in the genomes of both *P. syringae* CC1466 and CC1557

Similarity searches of the *P. syringae* genomes demonstrated that *hopM1*, which encodes an effector protein largely conserved throughout *P. syringae*, had likely been lost within strains CC1466 and CC1557. Further investigation demonstrated that an unknown protein open reading frame had replaced *hopM1* within these genomes (Figure 2). In order to test if this unknown protein is translocated into plant cells, we cloned the region encoding the predicted ORF into the translocation test vector pBAV178 and tested for delivery of AvrRpt2 as a fusion protein. We found that the resulting HopBJ1-AvrRpt2 fusion protein triggered cell death in both *Arabidopsis thaliana* Col-0 wild type and *rps2* lines, indicating that HopBJ1 itself was capable of triggering cell death in the absence of recognition of AvrRpt2 (Table 3, Figure 4B). In order to verify that HopBJ1 triggered cell death without AvrRpt2 we delivered HopBJ1:HA into a variety of plant genotypes using *Pto* DC3000 D28E (Cunnac et al., 2011). Delivery of a HopBJ1:HA can cause tissue collapse in a variety of Arabidopsis accessions (Table 3). All genotypes tested displayed strong cell death by either 24 or 48hr post inoculation.

**Table 3.** Responses of Arabidopsis Accessions to HopBJ1.

| <u>Arabidopsis Accession</u> | <u>Reaction</u> |
| --- | --- |
| Col | ++ |
| Col rps2 | ++ |
| Ws | ++ |
| Ws eds1 | ++ |
| Ws pad4 | ++ |
| Ws rar1 | ++ |
| Ws NahG | ++ |
| Ag-0 | ++ |
| Bur-0 | + |
| Can-0 | + |
| Ct-1 | ++ |
| Edi-0 | ++ |
| Hi-0 | + |
| Kn-0 | + |
| Ler-0 | ++ |
| Mt-0 | ++ |
| No-0 | ++ |
| Oy-0 | ++ |
| Po-0 | ++ |
| Rsch | ++ |
| Sf-0 | + |
| Tsu-0 | + |
| Wil-2 | + |
| Ws-0 | + |
| Wu-0 | + |
| Zu-0 | + |

++ HR in 24hr + HR in 48hr Gene names next to accession indicate mutant lines

**Figure 4.**
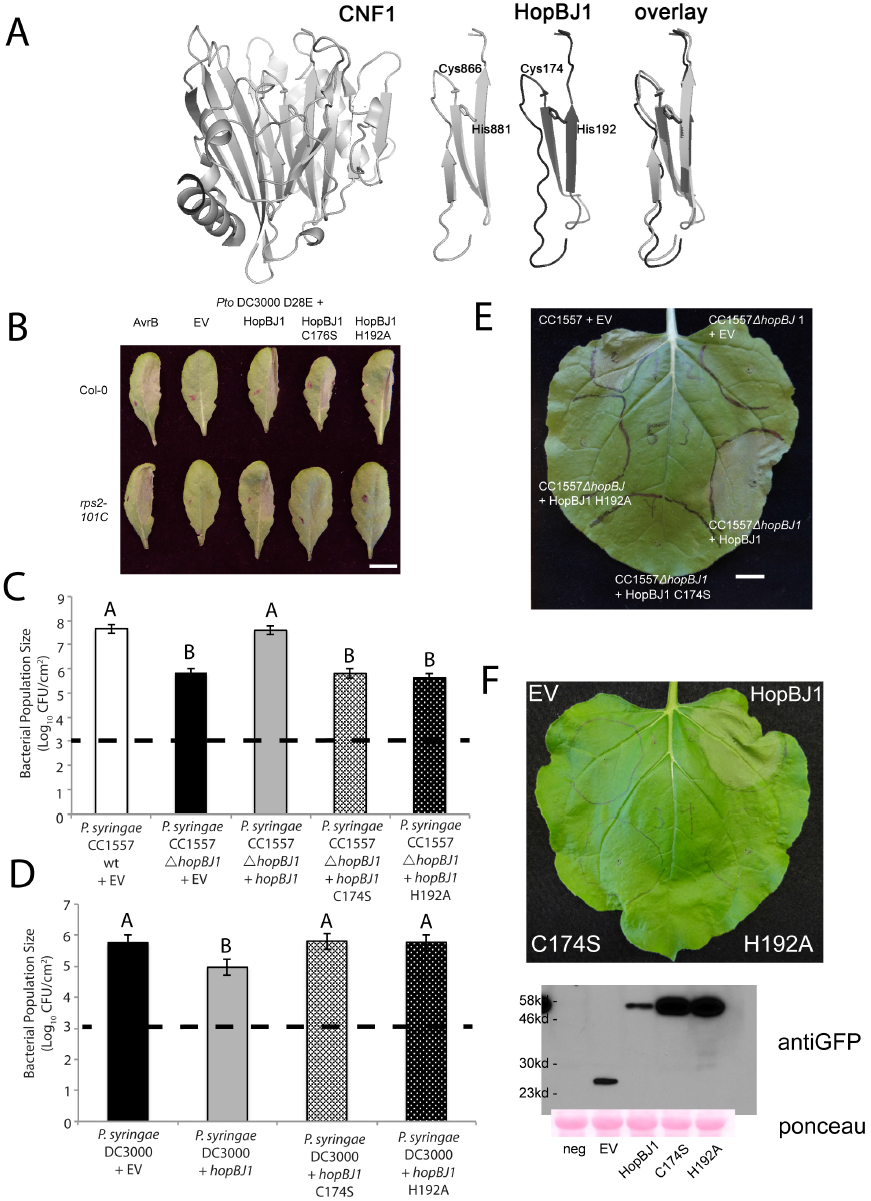
Protein Motifs and catalytic residues within HopBJ1 are similar to the *E. coli* CNF1 Toxin. **A)** Despite no sequence similarity between HopBJ1 and CNF1 by BLASTp searches, modeling of tertiary protein structure demonstrates limited similarity in protein structure between the two. Importantly, the catalytic dyad of *E. coli* CNF1 (Cys866-His881) perfectly lines up with a putative catalytic dyad within HopBJ1 (Cys174 and His192). **B)** All constructs of HopBJ1 can be translocated by *Pto* DC3000 D28E (Cunnac et al., 2011) into *Arabidopsis*. Top: translocation tests into accession Col-0. Bottom: translocation tests into *rps2*-101C mutant plants. Translocation of AvrB is shown as a positive control and causes HR in both plant backgrounds, while the *Pto* DC3000 D28E mutant causes no cell death. HopBJ1 causes rapid cell death across both plant backrounds when translationally fused to the C-terminus of AvrRpt2, while catalytic mutants only show cell death in Col-0. Picture is representative of >4 independent leaves per strain / plant combination **C)** Catalytic mutants of HopBJ1 cannot complement the growth defect of a CC1557 *hopBJ1* deletion strain. **D)** Translocation of HopBJ1, but not either of the two catalytic mutants, lowers growth of *Pto* DC3000 on *Arabidopsis* Col-0. **E)** Wild type and the complemented Δ*hopBJ1* deletion of strain CC1557 cause tissue collapse in *N. benthamiana*. Neither of the catalytic mutants complements the loss of virulence of the Δ*hopBJ1* deletion strain. Picture is representative of multiple assays. **F)** Transient expression of HopBJ1, but not the catalytic mutants, by *Agrobacterium* in *N. benthamiana* causes cell death. Western blots and Ponceau staining for each of these *Agrobacterium* infiltrations are displayed below. Picture is representative of multiple assays. Each growth curve experiment consisted of at least two independent trials, but representative data from only one trial is shown in each case. Letters indicate significant differences between bacterial population sizes between strains (p < 0.05, Tukey’s HSD). Dashed lines indicate approximate population sizes at day 0, which were not significantly different.

HopBJ1 induced cell death is not dependent on the known R-gene pathways as tissue collapse still occurs in *eds1*, *pad4* or *rar1* mutants (Table 3). HopBJ1-induced cell death also occurs in plants expressing the salicylic acid degrading enzyme NahG (Table 3). However, this rapid tissue collapse fails to occur out of a *hrcC-* background of *Pto* DC3000, indicating HopBJ1 secretion is dependent on a functioning T3SS (data not shown). The genomic region immediately upstream of *hopBJ1* possesses a canonical hrp-box sequence (Fig. S4), and so regulation of this gene very likely takes place through the action of the sigma factor HrpL (Mucyn et al., 2014). We therefore conclude that this novel ORF encodes a new putative T3E family, HopBJ, and that this family possesses broad cytotoxic capabilities.

#### Deletion of HopBJ1 from *P. syringae* CC1557 causes loss of disease symptoms and lowers bacterial populations *in planta*

We have created a deletion of *hopBJ1* in *P. syringae* strain CC1557, and have performed growth curves *in planta* using syringe infiltration. Deletion of *hopBJ1* causes a repeatable loss of growth *in planta* after two days of infection (Figure 3A). We further find that deletion of *hopBJ1* leads to a loss of the tissue collapse phenotype after 3 days (Figure 3B). We have been able to complement both of these phenotypes *in planta* through expression of *hopBJ1* using its native promoter *in trans* (Figure 3).

#### HopBJ1 resembles cytotoxic genes present within other species

Currently, the closest BLASTp hits for HopBJ1 are hypothetical proteins from *Serratia marcescens* (34% identity, YP_007346200.1) and *Hahella chejuensis* (30% identity, YP_435189.1). However, modeling of protein structure using the Phyre2 web server demonstrates that HopBJ1 displays limited similarity to the *E. coli* CNF1 toxin (Figure 4A). Surprisingly, amino acids critical for the deamidase functions of CNF1 appear to be conserved in HopBJ1 (Figure 4A) as well as in similar proteins from *S. marcescens* and *H. chejuensis* (Buetow et. al., 2001; data not shown). This modeling of protein structure further suggests that C174 and H192 are a functional catalytic dyad within HopBJ1.

#### Predicted catalytic residues of HopBJ1 are required for effector function

To test for functionality of predicted catalytic residues for HopBJ1, we measured the effect of putative catalytic dead mutants (C174S, H192A) *in planta*. Although mutant versions of HopBJ1 are successfully translocated by strain *Pto* DC3000 D28E into Arabidopsis, only the wild type version causes tissue collapse in *rps2* 101c plants (Figure 4B). Moreover, both catalytic dead mutants are unable to complement virulence defects of a *hopBJ1* deletion in CC1557 in *N. benthamiana* (Figures 5C and 5E). Both mutants, unlike the wild type version, also fail to lower the growth of *Pto*DC3000 in *Arabidopsis* (Figure 4D) and fail to cause tissue collapse when transiently expressed in *N. benthamiana.* Therefore, as with similar residues in *E. coli* CNF1 (Buetow et. al., 2001), we conclude that both C174 and H192 are required for full enzymatic function of HopBJ1.

## Discussion

Recent advances in genome sequencing have spurred great interest in conservation and diversification of virulence pathways across *Pseudomonas syringae* (O’Brien et al., 2011; Studholme, 2011). While genome gazing often leads to the identification of trends across a species, single observations must be treated with skepticism in the absence of additional data. One particularly interesting, yet relatively little understood, pattern across genomes involves a negative correlation in MLSA group II between size of T3E repertoire and presence of NRP pathways (Baltrus et al., 2011; Figure S1). Importantly, this trend does not appear to be a result of sampling bias or failure to discover novel T3E because strains within this clade have been thoroughly screened at both genetic and transcriptome level (Baltrus et al., 2011; Mucyn et al., 2014). Indeed the entire *hrpL* regulon, the major transcriptional regulator for all *P. syringae* T3E, is reduced within MLSA group II compared to other clades (Mucyn et al., 2014).

Strains CC1466 and CC1557 diverge early in the phylogeny of *P. syringae sensu latu*, and have not been isolated from diseased plants in nature (Morris et al., 2008). We have shown that CC1557 can act as a pathogen of *N. benthamiana* and, as with all other phytopathogenic *P.syringae* strains, that virulence requires a functional T3SS (Figure 1). We have found that both of these genomes contain few T3Es, with only 5 known loci present and shared by both (Table 2). Genomic comparisons between these two strains have further demonstrated that both contain an NRP toxin pathway very similar to those encoding syringomycin. As is the nature of NRP toxins even a single amino acid change can have dramatic consequences on the final toxin product chemistry (Bender et al., 1999), but it is important to note that these many of the predicted ORFs in these regions are highly similar (>90% protein ID) to syringomycin ORFs present within MLSA group II strains. Comparison to closely related outgroup strains demonstrates that acquisition of this NRP pathway is evolutionarily correlated with loss of T3E families. Indeed, the most closely related sequenced outgroup strain, CC1583, lacks this NRP pathway and contains twice the number of known T3E loci (Table 2). As such, we interpret these patterns as independent evidence suggesting an evolutionary or ecological tradeoff between presence of syringomycin and number of T3Es within the strain. We also note that there appears to be an additional, uncharacterized NRP pathway downstream of the syringomycin-like region within CC1557 (data not shown).

What selective pressure could underlie such a negative correlation between T3E and NRP toxins? It is possible that the functions of these NRP toxins are redundant with a specific suite of T3Es. In this case, T3E loss can take place because there is no longer positive selection to counter selection against evolutionary (such as recognition by the plant immune system) or physiological costs. For MLSA clade II and CC1557, acquisition of syringomycin and potentially additional NRP pathways could render the functions of some T3E obsolete. If this is the case, we further predict that the potential second NRP toxin pathway (downstream of the syringomycin-like pathway in CC1557) could have similar functions as syringopeptin. It is also possible that disease ecologies of most MLSA group II strains, as well as CC1466 and CC1557, differ compared to other focal *P. syringae* strains. We have a rudimentary understanding of differences in disease ecology throughout *P. syringae* pathogens, although it is recognized that differences do exist across the phylogeny (Clarke et al., 2010). Specifically, one clade of strains from MLSA group II has exchanged the canonical *P. syringae* T3SS with a second divergent T3SS at an independent genomic location while still maintaining toxin pathways. While these strains can grow *in planta*, it is unknown whether they can cause disease (Clarke et al. 2010). Furthermore, infection of woody hosts has arisen multiple independent times on multiple hosts yet such strains are still dependent on a functioning T3SS (O’Brien et al., 2011; Ramos et al., 2012). Strains that contain syringopeptin and syringomycin potentially cause disease on a wider host range than other groups, which may be enabled by generality of NRP toxins (Baltrus et al., 2011; Hwang et al., 2005; Quigley and Gross, 1994). Alternatively, these strains may survive and persist across a variety of environments and host plants, but may only rarely reach high enough population densities to cause disease. MLSA group II strains appear to survive better as epiphytes compared to other groups, and presence of NRP toxins may tie into this strategy (Feil et al., 2005). Moreover, such differences in disease ecology may be difficult to precisely measure because they may only be visible under natural conditions of infection and dispersal.

It is possible that there are a number of undescribed T3E families within CC1466 and CC1557, and to this point we have successfully used comparisons of genome synteny to identify HopBJ1. *hopBJ1* is present within the conserved effector locus within these strains, and as such is proximate to both the structural genes for the T3SS and the T3E *avrE*. While it is difficult to precisely map out the evolutionary history of this region, it does appear that *hopBJ1* has been recombined into this locus in place of *hopM1* (Figure 2). In some respects, recombination of *hopM1* from this locus is analogous to a situation witnessed in MLSA group I *P. syringae* strains where *hopM1* alleles have been cleanly swapped through recombination (Baltrus et al., 2011). HopM1 is an effector whose presence (although not necessarily in functional form) is conserved throughout *P. syringae sensu strictu*, and has been shown to act redundantly with AvrE during infection of plant hosts (Badel et al., 2006; Kvitko et al., 2009). Moreover, in contrast to frameshift or nonsense mutations disrupting the coding sequence of *hopM1* in *P. syringae*, presence of this gene is polymorphic within *P. viridiflava*. While some strains maintain an intact version, others lack *hopM1* completely (Araki et al.,2006; Bartoli et. al, 2014). It is currently unknown whether HopBJ1 can act redundantly with AvrE on specific hosts, although that our single deletion mutant shows a virulence phenotype on *N. benthamiana* speaks against this possibility. Moreover, since HopBJ1 appears to be responsible for a significant portion the growth of CC1557 *in planta*, it will be interesting to see how this effector behaves within different strain backgrounds on different hosts.

HopBJ1 itself displays protein structure similarity to CNF1 toxin found within pathogenic *E. coli* strains. CNF1 functions by causing deamidation of a glutamine residue, crucial for GTP hydrolysis for small GTPases of the Rho family, thereby leading to constitutive activation and actin disruption (Lemonnier et al., 2007). Furthermore, changing either of these amino acids within HopBJ1 eliminates virulence activity of CC1557 *in planta*, as well as during transient expression by *Agrobacterium*, and eliminates avirulence activity when delivered from *Pto*DC3000 in *Arabidopsis* (Figure 4). Potential functional parallels between CNF1 and HopBJ1 are particularly interesting because Rho GTPases are known to regulate cytoskeletal dynamics in plants (Mucha et al., 2011), actin has a demonstrated role in basal defense (Henty-Ridilla et al., 2013), and the *P. syringae* effector HopZ1a has been shown to target tubulin in order to promote virulence (Lee et al., 2012). If HopBJ1 does indeed target Rho family GTPases to disrupt the cytoskeleton, it would represent a striking example of molecular convergence across plant and animal pathogens.

The pathology of HopBJ1 on Arabidopsis, when delivered from strain *Pto* DC3000, suggests that HopBJ1 could be acting as a general toxin because cell death appears regardless of accession and is not dependent on typical R-gene related host defense pathways (Table 3). Moreover, HopBJ1 does act as an avirulence factor within *Pto*DC3000 because this cell death limits the growth of this strain in Arabidopsis (Figure 4D). In some ways, this may be similar to the functions of AvrE, which was identified as causing cell death in soybean (Kobayashi et al., 1989). It is tempting to think that both AvrE and HopBJ1 ultimately lead to the same physiological outcomes during infection, but so far little is known about processes targeted by AvrE.

What advantages HopBJ1 provides to CC1557 in nature remains unknown, as it is currently unclear whether this strain is capable of causing disease in any organism outside of the lab environment or syringe infiltration. Indeed, both CC1557 (snow) and CC1466 (asymptomatic plants) were originally isolated in order to characterize environmental diversity of *P. syringae* strains outside of crop disease. It is certainly possible that CC1557 and related strains persist at low levels across a range of plant hosts and habitats, using a T3SS and toxins to survive epiphytically but never reaching large enough population sizes to cause disease. It is also possible that CC1557 is naturally pathogenic within a suite of under-studied plant hosts or only under specific environmental conditions. However, given the disease symptoms measured under laboratory conditions, it is clear that CC1557 has the potential to cause rapid cell death in plant leaves relative to other *P. syringae* strains and that a significant amount of this activity is dependent on HopBJ1. These results suggest an intriguing possibility that the disease ecology of CC1557 is fundamentally different (for instance, more necrotrophic) than other commonly studied *P. syringae* strains.

Herein we have analyzed the genomes of two *P. syringae* strains isolated from environmental sources. These genomes contain a paucity of known T3E, even though virulence of CC1557 on *N. benthamiana* is dependent on a functioning T3SS. That these strains have also acquired a NRP pathway similar to syringomycin represents an independent evolutionary example that supports a negative correlation between NRP pathways and T3E repertoires. Furthermore, this data set highlights how genomic scans can lead to insights into pathogenesis while also demonstrating the power of genetic diversity to uncover genome-wide changes in virulence gene architecture.

## Methods

### Plasmids, bacterial isolates, and growth conditions

All bacterial strains and plasmids used or created are listed in table 1. Typically, *P. syringae and Agrobacterium* isolates were grown at 27°C on KB media using 50**μ**g/mL rifampicin. When necessary, cultures of *P. syringae, Agrobacterium* and *E. coli* were supplemented with antibiotics or sugars in the following concentrations: 10**μ**g/mL tetracycline, 50**μ**g/mL kanamycin, 50ug/ml spectinomycin, 25**μ**g/mL gentamycin (50ug/mL for Agrobacterium), and 5% sucrose.

All clones were created by first amplifying target sequences using *Pfx* polymerase (Invitrogen), followed by recombination of these fragments into the entry vector pDONR207 using BP clonase (Invitrogen). Site-directed mutagenesis was performed using SLIM amplification (Chiu et al., 2004). All ORF (without a *hrp*-box promoter) and gene (including the promoter) sequences were confirmed by Sanger sequencing of these pDONR207 clones. Clones in entry vectors were recombined into destination vectors of interest using LR clonase (Invitrogen).

### Genome Sequences and Searches

Draft genome assemblies for CC1466, CC1557,and CC1583 are publicly available through Genbank (accession numbers AVEM00000000, AVEH00000000, and AVEG00000000 respectively; Baltrus et al., 2014). One PacBio SMRT cell yielded 37,509 reads for a total of 268,122,626 nucleotides after filtering for quality. To piece together the complete genome sequence for CC1557, PacBio reads were first assembled using the HGAP software (Chin et al., 2013), yielding two contigs total with no scaffolding gaps. Contigs from the previous assembly (using only 100bp Illumina paired end reads, Baltrus et al. 2014) were then overlayed onto this HGAP assembly. When there was coverage by contigs from the Illumina assembly, the previous assembly sequence was chosen as the final sequence, but when there was no coverage the PacBio assembly was chosen. In cases where multiple divergent Illumina contigs assembled to the same region in the HGAP assembly, the Illumina sequence which matched the PacBio assembly was chosen. Since the second contig from the HGAP assembly possesses numerous plasmid related genes, we are confident that this does actually represent a plasmid present within this strain. Both the chromosome and plasmid were annotated using PGAAP (Angiuoli et al., 2008).

T3Es and NRP pathways were identified as per Baltrus et al. (2011). Briefly, draft genome assemblies were queried using tBLASTn with queries consisting of protein sequences for known type III effector families or for key loci within toxin pathways. Each BLAST hit was validated by hand for copy number, identity, and completeness. All BLAST results were visually inspected to make sure that each sequence displayed high similarity to only one region in the assembly, that this similar region was not part of a larger ORF, and that the length of this region was greater than 40% of the length of the original query sequence.

The Phyre2 server (http://www.sbg.bio.ic.ac.uk/phyre2/html/page.cgi?id=index) was used to carry out protein structure similarity searches (Kelley and Sternberg, 2009). In brief, this server performs an automated search of both the protein sequence and predicted folds for proteins of interest (in this case HopBJ1) across multiple databases. Within this pipeline, iterative searches such as PSI-Blast enable identification of distantly similar sequence matches. Once a profile is constructed for the sequence of interest, it is compared against a database of known protein structures and returns the best matches as a prediction of protein folds within the query.

### Generation of bacterial mutants

Bacterial mutants were generated as per Baltrus et al. 2012. Regions (>500bp) upstream and downstream of the target genes were PCR amplified separately and then combined into one fragment by overlap extension PCR. The bridge PCR amplicon was then cloned into pDONR207, and further moved into pMTN1907 using LR clonase. Once mated into *P. syringae*, single recombination of a homologous region upstream or downstream of the target region and subsequent selection on sucrose allows for screening of clean deletions. Mutants were confirmed by phenotyping for sucrose resistance, tetracycline sensitivity, PCR amplification of the deletion, and failure of PCR to amplify regions within the deletion.

### Negative Correlation Between Syringomycin-like Toxin Pathways and Type III Effectors

We calculated statistical significance for negative correlation between toxin pathways and effectors using data on the number of full length T3Es per strain from Baltrus et al. 2011 as well as data from Table 2 (Figure S2). To minimize bias due to phylogenetic relationships, we chose > 5 diverse strains from MLSA groups 1, 2, and 3 as well as CC1557 and CC1583. We then performed a Wilcoxian rank sum test to compare the number of full-length effectors between genomes which either contain or lack the genetic capacity for production of a syringomycin-like toxin.

### In planta Growth Curves, Cell-death assays and Translocation Tests

*Nicotiana benthamiana* plants were grown for 4-6 weeks on a long day cycle (16 hours light/8 hours dark). Plants were removed from the growth chamber and allowed to acclimate to ambient laboratory conditions for a period of 3-5 days prior to infiltration (Arizona). Bacteria were cultured over night on KB amended with proper antibiotics. Bacterial cells were washed 1x in and resuspended to an OD_600_ of 0.002 in sterile 10 mM MgCl_2_ yielding a final inoculum density of approximately 1×10^5^ (North Carolina) or 1×10^6^ CFU/mL (Arizona). Cell suspensions were then syringe infiltrated into the abaxial side of *N. benthamiana* leaves. Populations were recovered after 2 (North Carolina) or 3 (Arizona) days of growth using a corkborer unless otherwise noted. Leaf disks were disrupted using glass beads and a bead-beater device and populations were enumerated by dilution plating onto KB amended with appropriate antibiotics. Arabidopsis growth assays were performed similarly, with the exception that plants were grown in walk-in rooms maintained at on a short day cycle (9 hours light at 21C and 15 hours dark at 18C).

Cell death induced by hopBJ1 was assayed after transient expression of 35S-driven hopBJ1:YFP in *N. benthamiana.* Bacteria were grown to saturation overnight in 2xYT media and then diluted to an OD_600_ of 0.1. Cultures were syringe injected into fully expanded leaves of 5-6 week old plants and cell death was visualized 24hr post-innoculation. Accumulation of WT and mutant HopBJ1 was verified with standard Western blotting of lysates from injected leaf cores using anti-GFP (Roche)

HopBJ1 was tested for the ability to secrete the active C-terminal fragment of AvrRpt2 to cause a hypersensitive response (HR) in Arabidopsis accession Col-0 (Guttman et al., 2002). For this test, promoterless *hopBJ1* was cloned into plasmid pBAV178 (Vinatzer et al., 2006) so that expression in strain *Pto* DC3000 D28E was dependent on a *tet* promoter. This construct was also placed into a *hrcC-* mutant of *Pto* DC3000 (Baltrus et al., 2012), in order to test the requirement of a functioning TTSS for tissue collapse. All HR tests were performed on ∼5 week old plants and utilized a bacterial density of OD_600_ 0.05. Tissue collapse was measured at either 24 or 48 hours post infection and confirmed by at least four independent tests. For translocation of hopBJ1 and hopBJ1 site-directed mutants, similar assays were done with native promoter clones in pJC532 delivered from the D28E effector-deleted strain of *Pto* DC3000. (Chang et al., 2005)

### Phylogenetic Methods

Phylogenetic analyses were performed on concatenated MLSA loci as described previously using concatenated fragments from 7 MLSA genes across a diverse array of isolates from P. syringae (Baltrus et al, 2011) as well as AvrE and HrpW protein sequences. MrBayes was used to perform Bayesian phylogenetic analyses with flat priors, a burn-in period of 250,000 generations, and convergence after 1,000,000 total generations (Ronquist et al., 2012).

## Acknowledgements

Funding was provided by startup funds to David Baltrus from the University of Arizona. We thank the University of Delaware Sequencing Core Facility for technical help with PacBio sequences. We thank Cindy Morris, INRA, France for kindly providing access to strains described in this manuscript. We also thank Jeff Dangl for experimental reagents and positive encouragement.

---

**Figure S1.**
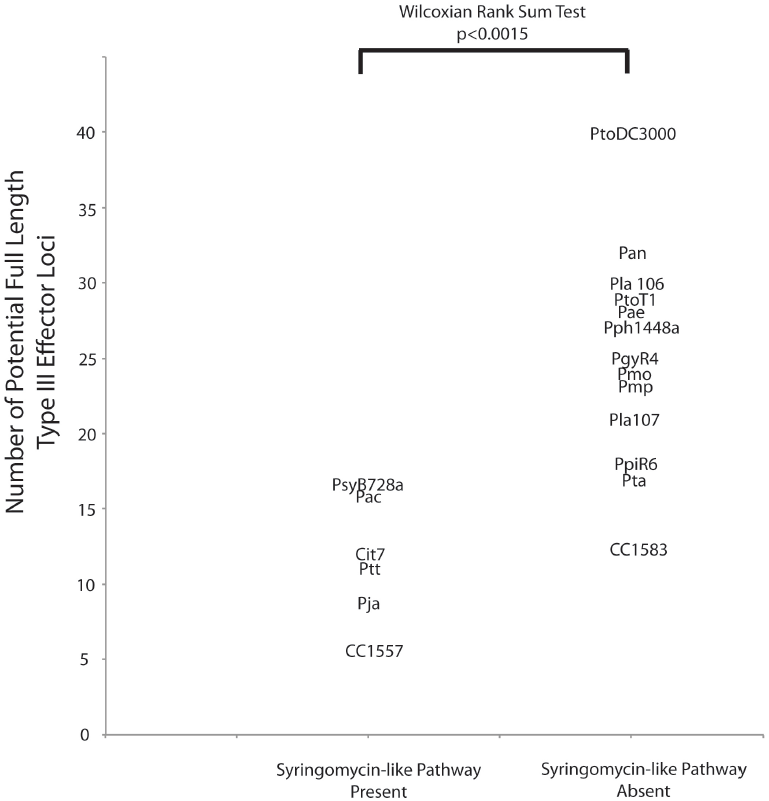
Negative Correlation Between Type III Effector Number and Presence of Syringolin Pathways. The number of type III effectors is plotted for a subset of strains investigated within Baltrus et al. 2011 as well as CC1557 and CC1583, with strains grouped by the presence of a syringomycin-like pathway. Full strain names are described in Baltrus et al. 2011. Genomes with the genetic potential to encode a syringomycin-like toxin contain a significantly lower number of type III effectors (Wilcoxian rank sum test, p<0.0015).

**Figure S2.**
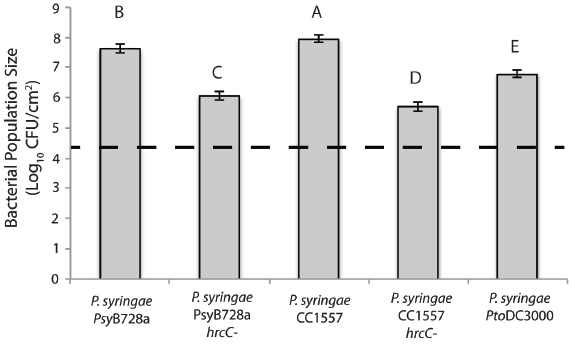
*P. syringae* CC1557 grows as well as *P. syringae* pv. *syringae* B728a on *N. benthamiana*. Bacterial population sizes after one day of growth are shown for wild type and *hrcC-* versions of the established *N. benthamiana* pathogen *P. syringae* pv. *syringae* B728a. Also shown are population sizes for wild type and *hrcC-* versions of strain *CC1557* as well as wild type *P. syringae* pv. *tomato* DC3000. Dashed line indicates approximate day 0 population sizes for all strains, which were not significantly different than one another. Error bars represent 2 standard errors. Growth curves were performed twice with similar results, but only data from one trial are shown. Letters indicate significant differences in population sizes between strains (p<0.05, Tukey’s HSD)

**Figure S3.**
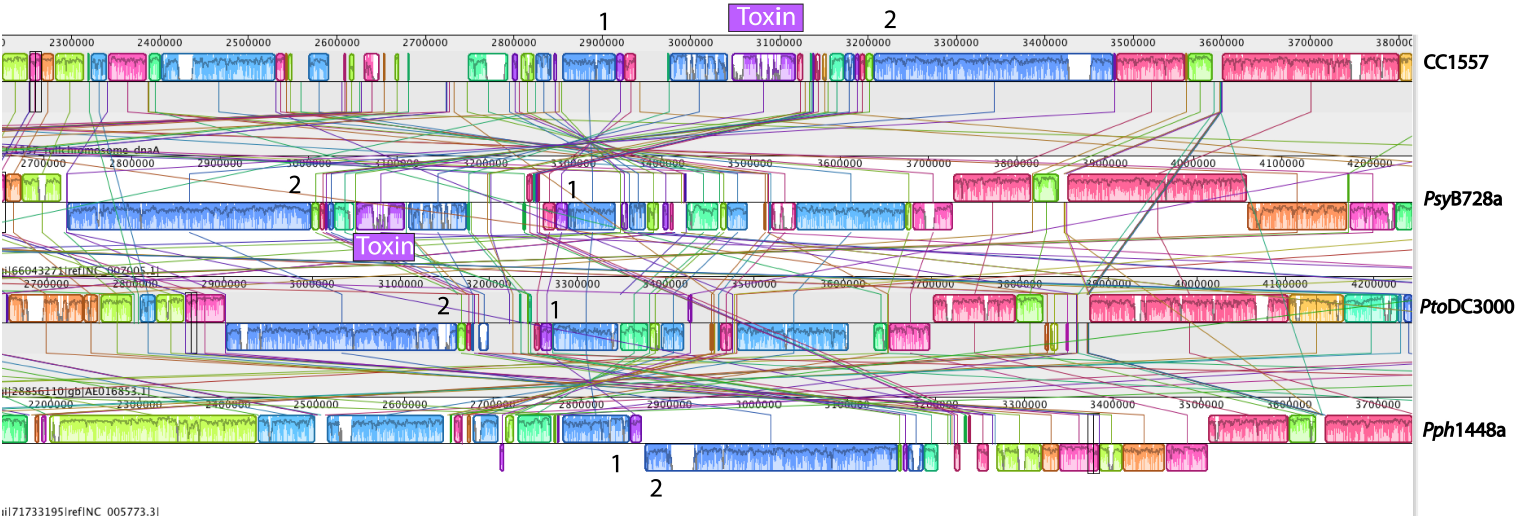
Toxin Regions are Syntenic in *P. syringae* pv. *syringae* B728a and *P. syringae* CC1557. Mauve was used to align four whole *P. syringae* genomes, and the region that contains pathways encoding syringomycin was extracted from *P. syringae* pv. *syringae* B728a. The toxin region for syringomycin and syringopeptin is shown in purple, while adjacent genomic regions conserved throughout the four genomes are listed with either the number 1 or 2. The strain order is (from top): CC1557, pv. *Syringae* B728a, pv. *phaseolicola* 1448a, pv. *tomato* DC3000

**Figure S4.**
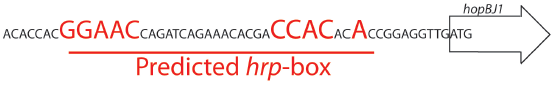
Putative *hrp*-box for *hopBJ1* in *P. syringae* CC1557. The sequence of the coding strand around *hopBJ1* is shown from strain CC1557. The ORF of *hopBJ1* is shown as an arrow. The region immediately upstream the *hopBJ1* start codon is also shown, with the predicted *hrp-*box and consensus nucleotides labeled in red.

